# 3D genome organization around nuclear speckles drives mRNA splicing efficiency

**DOI:** 10.1101/2023.01.04.522632

**Authors:** Prashant Bhat, Amy Chow, Benjamin Emert, Olivia Ettlin, Sofia A. Quinodoz, Yodai Takei, Wesley Huang, Mario R. Blanco, Mitchell Guttman

## Abstract

The nucleus is highly organized such that factors involved in transcription and processing of distinct classes of RNA are organized within specific nuclear bodies. One such nuclear body is the nuclear speckle, which is defined by high concentrations of protein and non-coding RNA regulators of pre-mRNA splicing. What functional role, if any, speckles might play in the process of mRNA splicing remains unknown. Here we show that genes localized near nuclear speckles display higher spliceosome concentrations, increased spliceosome binding to their pre-mRNAs, and higher co-transcriptional splicing levels relative to genes that are located farther from nuclear speckles. We show that directed recruitment of a pre-mRNA to nuclear speckles is sufficient to drive increased mRNA splicing levels. Finally, we show that gene organization around nuclear speckles is highly dynamic with differential localization between cell types corresponding to differences in Pol II occupancy. Together, our results integrate the longstanding observations of nuclear speckles with the biochemistry of mRNA splicing and demonstrate a critical role for dynamic 3D spatial organization of genomic DNA in driving spliceosome concentrations and controlling the efficiency of mRNA splicing

## INTRODUCTION

The nucleus is highly organized such that DNA, RNA and protein molecules involved in transcription and processing of distinct RNA classes (e.g., ribosomal RNA, histone mRNAs, snRNAs, mRNAs) are spatially organized within or near specific nuclear bodies [1–5] (e.g., nucleolus [6,7], histone locus body [8,9], Cajal body [9–11], nuclear speckles [12,13]). Yet, despite being first described more than a century ago, the functional roles of these nuclear bodies remain untested [14–16]. In theory, they could represent structures that are critical for transcription and/or processing of specialized classes of RNA [2], or instead they could represent an emergent property of co-regulation whereby regions of shared regulation simply self-assemble in three-dimensional (3D) space [17]. Distinguishing between these possibilities has proven challenging [14–16] because many of the molecular components contained within these nuclear bodies serve dual roles – as catalytic components required for transcription or RNA processing and as structural components required for the integrity of these structures [18–22].

To explore this question, we focused on the relationship between nuclear structure and mRNA splicing. In higher eukaryotes, most RNA Polymerase II (Pol II) transcribed genes contain intronic sequences that must be removed from precursor messenger RNAs (pre-mRNAs) to generate mature mRNA transcripts [23,24]. mRNA splicing is predominantly co-transcriptional such that nascent pre-mRNAs are spliced as they are transcribed [25–31]. Incomplete splicing yields mRNAs that are degraded by nonsense-mediated decay and results in decreased protein levels [32], and disruption of mRNA splicing is associated with many human diseases [33] including cancer [34–36], neurodegeneration [37–40], and immune dysregulation [41,42]. Due to this central importance, splicing needs to be highly efficient to ensure the fidelity of mRNA and protein production.

Early studies visualizing the localization of mRNA splicing factors– including proteins (e.g., SRRM1, SRSF1, SF3a66) and non-coding RNAs (e.g., U1, U2) [43,44] – observed that these factors were not uniformly distributed throughout the nucleus but instead were enriched within specific, 3D territories referred to as nuclear speckles [45–47]. Because of the preferential localization of splicing regulators, nuclear speckles were initially thought to represent the site of mRNA splicing in the nucleus [12,13]. However, subsequent studies showed that splicing does not occur within nuclear speckles, but instead splicing factors diffuse away from speckles to bind nascent pre-mRNAs and catalyze the splicing reaction [48–52]. These observations led to the prevailing notion that nuclear speckles simply act as storage bodies of inactive spliceosomes rather than functional structures involved in mRNA splicing [53–58]. Accordingly, despite their initial description over 40 years ago [45–47], what functional role, if any, speckles might play in the process of mRNA splicing remains unknown [59].

Recently, we developed genome-wide methods to explore the higher-order three-dimensional organization of DNA and RNA in the nucleus [60–62]. Using these and related approaches [63,64], we and others identified that nuclear speckles represent major structural hubs that organize interchromosomal contacts corresponding to genomic regions containing highly transcribed Pol II genes and their associated nascent pre-mRNAs [61,62]. Because co-localizing splicing factors (enzymes) and their target pre-mRNAs (substrates) would concentrate splicing factors at the locations where they must act (nascent pre-mRNA), we hypothesized that organization of highly transcribed Pol II genes on the periphery of nuclear speckles would increase the concentration of spliceosomes at these nascent pre-mRNAs, thereby increasing their splicing efficiency. In this way, spatial organization may act to effectively couple Pol II transcription and mRNA splicing efficiency. Here we demonstrate an essential role for 3D organization of genomic DNA in controlling the efficiency of mRNA splicing.

## RESULTS

### snRNAs preferentially bind pre-mRNAs of genes that are close to speckles

To explore DNA localization around the nuclear speckle, we first computed speckle contacts for all genomic regions using both genomic (RNA & DNA SPRITE) [62] and microscopy (seqFISH+) [64] approaches in mouse embryonic stem (ES) cells. We observed that DNA regions that exhibit high SPRITE-based speckle contact frequencies (e.g., Tcf3, Foxj1, and Nrxn2) were preferentially located adjacent to SF3a66, a protein marker of nuclear speckles (Figure 1A). Conversely, DNA regions with low SPRITE-based speckle contact frequencies on the same chromosomes (e.g., Grik2, Efemp1, Zfand5) were located farther away from SF3a66 foci (Figure 1A). Comparing 2,460 paired genomic regions, we observed that SPRITE-based speckle contact frequency and DNA distance to SF3a66 were inversely correlated (r = −0.72), indicating that SPRITE accurately measures genomic distance to nuclear speckles (Figure 1B). We refer to genomic regions with the highest 5% of speckle contact frequencies as speckle close and those with the lowest 5% as speckle far.

**Figure 1:**
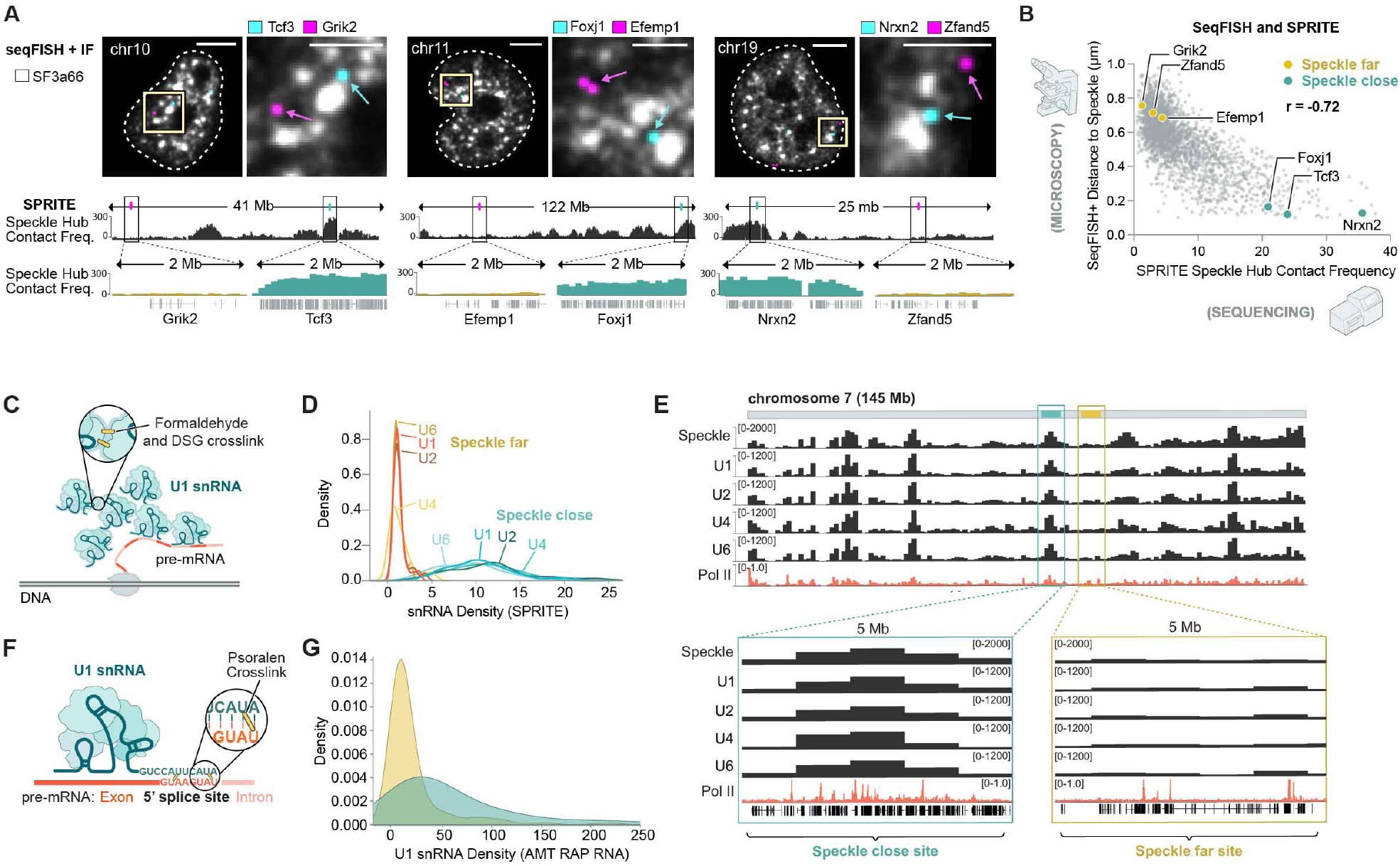
snRNAs preferentially bind pre-mRNAs of genes that are close to speckles. **(A)** Three reconstructed images for DNA seqFISH+ and immunofluorescence (SF3A66) in mouse ES cells comparing speckle close genes (Tcf3, Foxj1, Nrxn2 in blue) and speckle far genes (Grik2, Efemp1, Zfand5 in purple) (top). Images are maximum intensity z-projected for 1 μm section. White lines represent nuclear segmentation. Scale bars in zoom out panels are 5 μm and zoom in panels are 2.5 μm. Speckle contact frequencies from SPRITE for chromosomes 10, 11, and 19 at 100-kb resolution (bottom). Zoom in, speckle contact frequencies from SPRITE for the 2 Mb region around genes shown in top. **(B)** Genome-wide comparison of DNA seqFISH+ distance to exterior of speckle (μm) and SPRITE speckle hub contact frequency for 2460 paired genomic regions. Pearson r correlation is −0.72. **(C)** Schematic of types of RNA-DNA interactions captured by SPRITE. Formaldehyde and DSG crosslink nucleic acids and proteins to each other and SPRITE can measure the number, type (DNA or RNA), and sequence of molecules within each crosslinked complex. **(D)** Normalized density of U1, U2, U4, U6 snRNAs on speckle close versus speckle far genomic regions. Normalization for each snRNA is to the mode of the speckle far distribution to visualize all snRNA densities on the same scale. RPKM for both speckle far and close genes is thresholded between 2.5-7.5. **(E)** Whole chromosome 7 view of SPRITE contact frequencies at 1-Mb resolution for speckle hub, U1, U2, U4 and U6 snRNAs. Pol II-S2P ChIP-seq density at 100-kb resolution. **(F)** Schematic of direct RNA-RNA interactions capture by AMT RAP RNA67. Psoralen forms direct crosslinks between RNA-RNA hybrids, affinity purification (not shown) selectively captures U1 snRNA, and all directly hybridized pre-mRNAs. **(G)** U1 snRNA density from AMT RAP RNA for speckle close versus speckle far regions.

Having defined genome-wide proximity to nuclear speckles, we explored the localization of the spliceosome – the molecular machinery that carries out splicing and consists of U-rich small nuclear RNAs (snRNAs) and associated proteins [65] – across the genome. We considered two possible models for spliceosome association with pre-mRNA. In the direct-recruitment model, the spliceosome is directly recruited by either Pol II or the nascent pre-mRNA, which would result in the spliceosome associating with transcribed regions proportional to their mRNA levels. Alternatively, in the speckle-recruitment model, the spliceosome would accumulate preferentially at nascent pre-mRNAs that are localized near nuclear speckles.

To test these two models, we mapped the localization of the U1, U2, U4, and U6 snRNAs across the genome using RNA & DNA SPRITE (RD-SPRITE, Figure 1C). As expected, these snRNAs are enriched over genomic DNA regions that are actively transcribed into pre-mRNA. However, rather than simply reflecting pre-mRNA levels as would be predicted by the direct-recruitment model, we observed that regions that are close to nuclear speckles display ~10-fold higher enrichment of snRNAs independent of gene expression levels (Figure 1D, Supplemental Figure 1A-E). For example, two neighboring genomic regions on mouse chromosome 7 that are transcribed at comparable levels, but that are located within a speckle close and speckle far region display a ~4-fold difference in snRNA levels (Figure 1E). These results indicate that spliceosome concentrations are highest at nascent pre-mRNAs that are in proximity to nuclear speckles.

Because RD-SPRITE utilizes protein-protein crosslinking (formaldehyde + DSG) to map RNA-DNA contacts [60], this approach captures associations that are indirect and therefore may not reflect the proportion of pre-mRNAs directly engaged by spliceosomes [61,62] (Figure 1C). To measure the number of spliceosomes that directly bind to nascent pre-mRNAs, we used psoralen-mediated crosslinking (which forms covalent crosslinks only between directly hybridized nucleic acids [66]) to map U1 interactions with pre-mRNAs (Figure 1F). We previously showed that this approach is highly specific at mapping U1 binding to 5’ splice sites at exon-intron junctions [67]. Using this data, we computed the frequency of U1 binding to each pre-mRNA (number of U1 bound RNAs divided by RNA abundance) and compared U1 binding frequency to the distance between the nascent locus and nuclear speckles. We observed ~3-fold higher levels of U1 binding to pre-mRNAs transcribed from speckle close genes compared to those transcribed from speckle far genes (Figure 1G).

Together, these results indicate that proximity of genomic DNA regions to nuclear speckles is associated with increased concentrations of spliceosomes and spliceosome engagement on pre-mRNA.

### Co-transcriptional splicing efficiency varies based on proximity to nuclear speckles

Because the efficiency of a reaction is dependent on substrate and enzyme concentration, we reasoned that higher concentration of spliceosome components (enzyme) at pre-mRNAs (substrate) located proximal to nuclear speckles would lead to increased co-transcriptional splicing efficiencies (e.g., the proportion of spliced products to total mRNA produced, Figure 2A) relative to pre-mRNAs that are located farther from the speckle.

**Figure 2:**
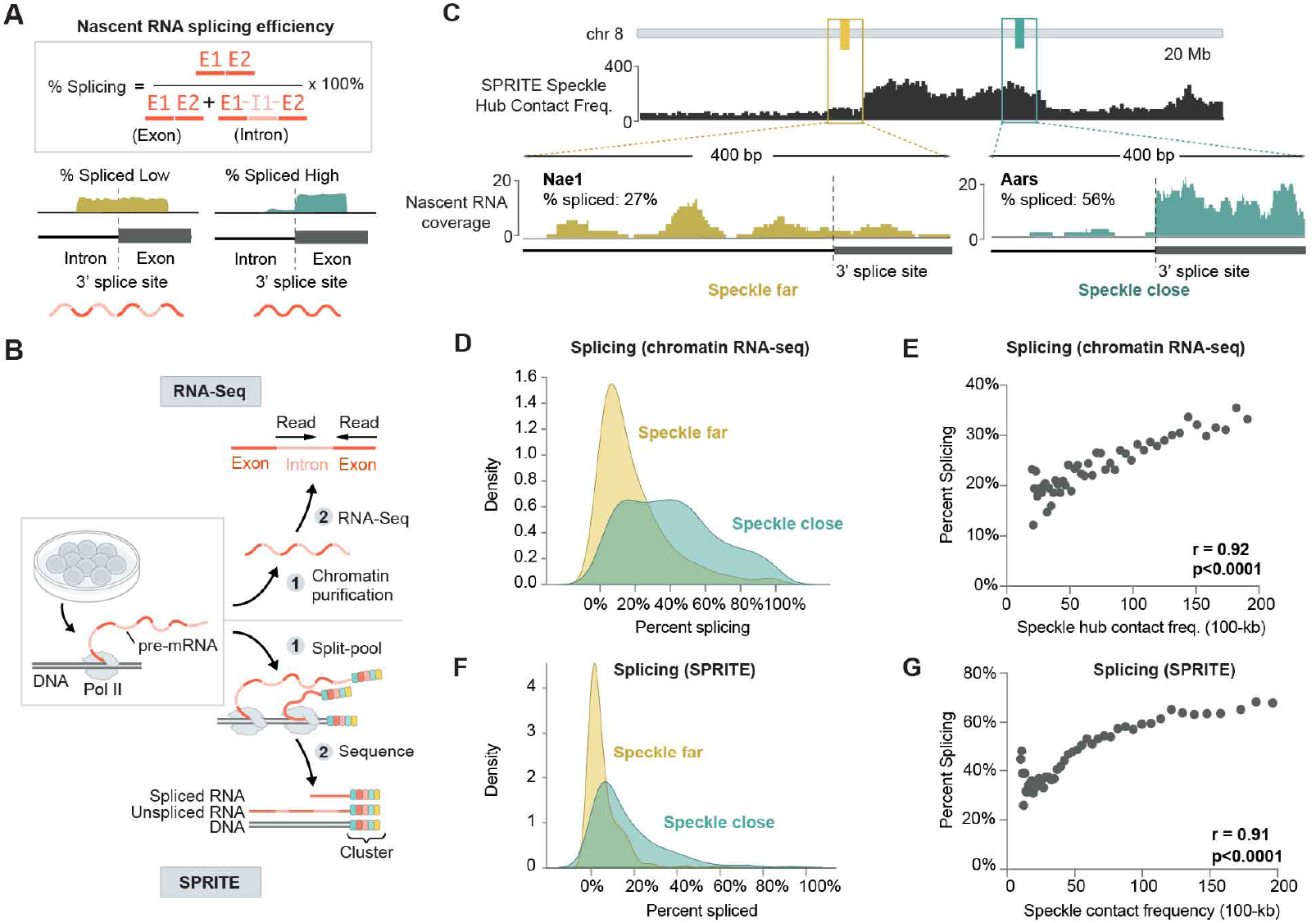
Co-transcriptional splicing efficiency varies based on proximity to nuclear speckles. **(A)** Nascent RNA splicing efficiency calculation. Splicing efficiency of a gene is calculated by taking the ratio of exon to total pre-mRNA counts from RNA sequencing (exons + introns). **(B)** Schematic of nascent RNA sequencing and SPRITE methods used to measure splicing efficiency. **(C)** SPRITE speckle hub contact frequency for a 20-Mb region on chromosome 8 (top). Nascent RNA coverage from chromatin RNA sequencing for a speckle far (Nae1) and speckle close (Aars) gene around a single 3’splice site (bottom). Percent spliced across entire gene is 27% (Nae1) and 56% (Aars). **(D)** Density plot of percent spliced for genes located within speckle close or speckle far 100-kb genomic regions (461 speckle close genes and 460 speckle far genes). **(E)** SPRITE speckle hub contact frequency (x axis) and percent spliced for genes from nascent RNA sequencing within each bin (y axis) across 50 bins. Each point/bin contains at least 20 genes and reflects the average splicing for that bin. Pearson r correlation = 0.92. **(F)** Density plot of percent spliced within 100-kb genomic intervals from SPRITE for speckle close and speckle far regions (312 speckle close and 311 speckle far 100-kb regions). **(G)** SPRITE speckle hub contact frequency (x axis) and percent spliced within genomic bins from SPRITE (y axis) across 50 bins. Each point/bin contains at least 20 regions and reflects the average splicing for that bin. Pearson r correlation = 0.91.

To focus on splicing of pre-mRNAs that occurs near the DNA locus from which it is transcribed (which we refer to as co-transcriptional splicing), we analyzed nascent RNA that is associated with chromatin using a stringent biochemical purification procedure [68,69] (Figure 2B). Using these data, we computed the splicing efficiency for each gene by taking the ratio of spliced reads relative to total pre-mRNA reads (spliced counts + unspliced counts) (Figure 2A). Overall, we observed that genes that were located closest to nuclear speckles showed a >2-fold higher splicing ratio compared to genes that are farthest from nuclear speckles (41.0% vs 19.1%) (Figure 2C-D). More generally, we observed a strong correlation between speckle contact frequency and splicing efficiency (r=0.92, p<0.0001, Figure 2E).

To further validate this effect and exclude the possibility that the observed splicing differences might reflect mature mRNA in our biochemical purification, we used an orthogonal method to measure mRNA levels on chromatin. Specifically, we used RD-SPRITE to analyze splicing ratios of RNAs [70] exclusively when they were associated with the DNA of their own nascent locus (Figure 2B). We then computed splicing efficiency as the fraction of exons over the total number of exons and introns. Consistent with the chromatin RNA-Seq data, we observed ~3 fold higher splicing in speckle-close (16.1%) to speckle-far (5.5%) regions (Figure 2F). Furthermore, we observed a strong correlation between the splicing efficiency per gene and its speckle contact frequency (r=0.91, p<0.0001; Figure 2G).

Together, these results indicate that the pre-mRNA splicing efficiency is highest for speckle-associated genes and that this splicing efficiency is achieved while the pre-mRNA is bound at its nascent locus.

### pre-mRNA organization around nuclear speckles is sufficient to drive increased mRNA splicing

Because genes differ in multiple ways beyond their nuclear speckle proximity (e.g., gene length, alternative splicing patterns, and sequence-specific features), it remains possible that the observed increase in splicing efficiency is due to other gene-specific or genomic DNA features (e.g., chromatin structure) that might also correlate with speckle proximity.

To directly test whether speckle proximity drives splicing efficiency, we designed a splicing reporter that can be directly recruited to nuclear speckles, allowing us to measure its splicing efficiency within individual cells. Specifically, we generated a reporter that produces an mRNA that is translated into GFP when spliced, but not when unspliced (Figure 3A). Increased GFP signal reflects increased reporter splicing and can be quantitatively measured within each cell via a fluorescence readout (Figure 3A). In the intron of this reporter, we embedded an MS2 bacteriophage RNA hairpin that binds with high affinity to the MS2 bacteriophage coat protein (MCP) [71]. We used this system to localize the pre-mRNA reporter to specific nuclear locations by co-expressing the splicing reporter together with specific MCP-fusion proteins that are known to localize at different locations within the nucleus (Figure 3B). Specifically, we expressed SRRM1 and SRSF1, two proteins that localize within nuclear speckles [22,72]. SRRM1 is primarily localized in nuclear speckles (punctate), while SRSF1 exhibits both speckle (punctate) and nucleoplasmic (diffuse) localization. As controls, we expressed several non-speckle proteins, including SRSF3 and SRSF9 (two splicing proteins that are not enriched within nuclear speckles but are localized throughout the nucleoplasm [73,74]) and LBR (a protein that is anchored in the nuclear membrane and associates with the transcriptionally inactive nuclear lamina [75]).

**Figure 3:**
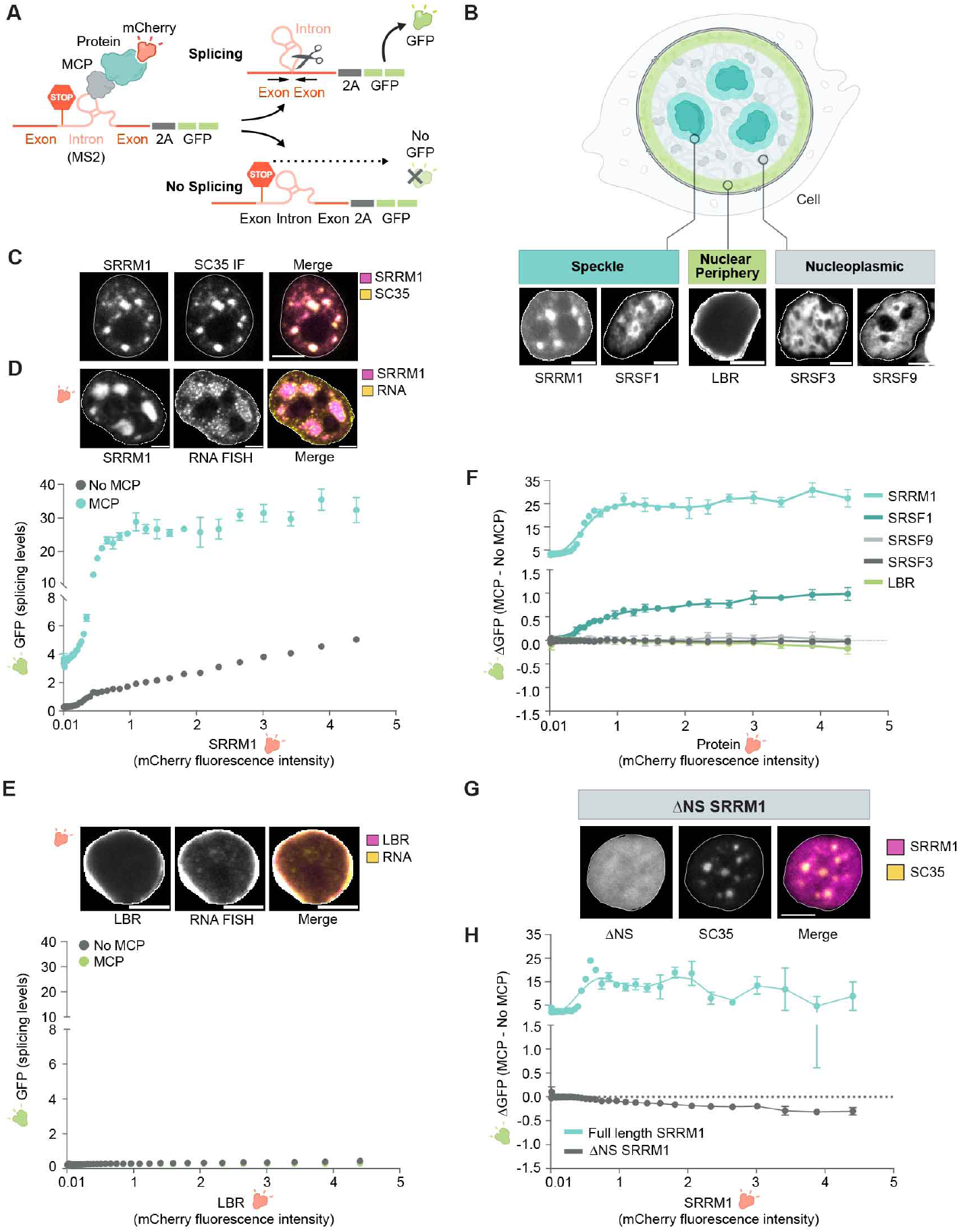
pre-mRNA organization around nuclear speckles drives splicing efficiency. **(A)** Schematic of pre-mRNA splicing assay via a fluorescence based read out. Individual proteins of interest are mCherry-tagged (shown) or without (not shown) an MCP tag. MCP protein binds to the complementary MS2 stem loop embedded within the intron of the pre-mRNA reporter. GFP is expressed only when the reporter is spliced and measured via FACS. **(B)** Schematic of specific nuclear locations (speckle, nuclear periphery, nucleoplasm, top) and mCherry fluorescence of their corresponding proteins (SRRM1, SRSF1; LBR; SRSF3, SRSF9, bottom). Nucleus is outlined in white. Scale bar is 5 μm. **(C)** Fluorescence microscopy for mCherry-SRRM1 (top left). co-immunofluorescence for SC35 (top middle), and merge (top right). Scale bar is 5 μm. **(D)** Localization of SRRM1+MCP with mCherry reporter and single-molecule RNA FISH. Nucleus is outlined in white. Scale bars, 5 μm (top). GFP levels (x axis) versus fluorescence intensity (levels) of SRRM1 (y axis) (bottom). Error bars are S.E.M for three replicates. **(E)** Localization of LBR+MCP with mCherry reporter and single-molecule RNA FISH. Nucleus is outlined in white. Scale bars, 5 μm (top). GFP levels (x axis) versus fluorescence intensity (levels) of LBR (y axis) (bottom). Error bars are S.E.M for three replicates. **(F)** Difference of GFP expression between constructs with MCP and no MCP (y axis) versus mCherry fluorescence intensity (x axis) for all constructs tested. Error bars are S.E.M for three replicates. **(G)** Fluorescence microscopy for mCherry-SRRM1-ΔNS (bottom left). co-immunofluorescence for SC35 (bottom middle), and merge (bottom right). Error bars are S.E.M for three replicates. Scale bar is 5 μm. **(H)** Difference of GFP expression between SRRM1 full length and SRRM1 ΔNS constructs with MCP and no MCP (y axis) versus mCherry fluorescence intensity (x axis). Error bars are S.E.M.

We transfected each of these proteins fused to MCP and mCherry (to directly visualize localization) and, using fluorescence microscopy, confirmed that each protein localized in the nucleus as expected (Figure 3B, Supplemental Figure 2A-E). We observed that SRRM1-MCP co-localized with endogenous SC35, a well-characterized marker of nuclear speckles (Figure 3C), while SRSF3 and SRSF9 localized diffusively throughout the nucleus and LBR localized to the periphery of the nucleus (Figure 3B, Supplemental Figure 2A-E). Next, we confirmed that the MS2-containing reporter RNA co-localized along with the MCP fusion protein using RNA FISH coupled with fluorescence microscopy of mCherry (Figure 3D–4E). We observed that the MS2-RNA localizes within nuclear speckles when co-expressed with SRRM1-MCP and localizes at the nuclear periphery when co-expressed with LBR-MCP. As expected, cells that express higher concentrations of the MCP-fusion protein exhibit increased co-localization of MS2-RNA (Supplemental Figure 3).

**Figure 4:**
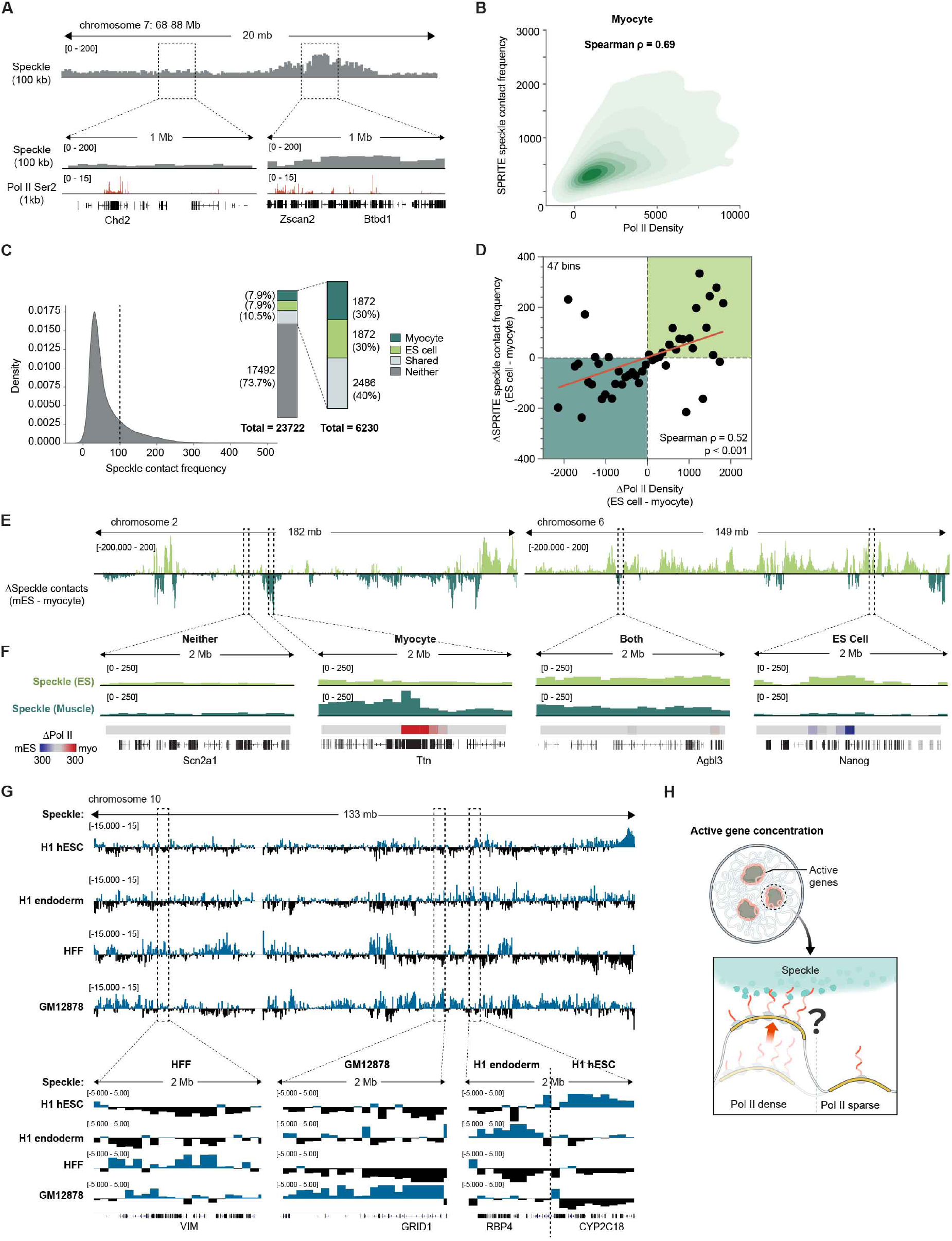
Differential gene positioning around nuclear speckles corresponds to differential Pol II occupancy. **(A)** SPRITE speckle hub contact frequency at 100-kb resolution for a 20-Mb region on chromosome 7 in mouse myocytes. Pol II-S2P ChIP-seq density at 1-kb resolution. **(B)** Ser2-P Pol II density (x axis) and normalized SPRITE speckle contact frequency (100-kb resolution) for myocytes. Spearman correlation = 0.69; p<0.0001. **(C)** Distribution of SPRITE speckle contact frequencies (100-kb resolution) for normalized mES and myocyte cell SPRITE (left). Distribution of number of genomic regions categorized as speckle hubs in myocyte, ES cells, both, or neither (right). **(D)** Difference in Ser2-P Pol II density (x axis) versus difference in SPRITE speckle hub contact frequency (y axis) between ES cells and myocytes at 1 Mb resolution. 47 bins. Spearman correlation = 0.52; p<0.001. **(E)** Difference in speckle hub contact frequency between mESCs (bottom) and myocytes for chromosomes 2 and 6. **(F)** 2-Mb zoom in regions of speckle contact frequencies and Ser2P Pol II densities for Scn2a1 (speckle in neither), Ttn (myocyte specific), Agbl3 (speckle in both) and Nanog (mES cell specific). **(G)** Difference in SPRITE speckle hub contact frequency for chromosome 10 between sample and average of the other three samples for each of H1 hESC, H1 endoderm, HFF, and GM12878 human cell lines. Zoom ins are 2-Mb regions of speckle contact frequencies for VIM (HFF specific), GRID1 (GM12878 specific), RBP4 (endoderm specific), and CYP2C18 (H1 hESC specific). **(H)** Model explaining how Pol II density may act to reposition genomic DNA into proximity with the nuclear speckle.

Having demonstrated the ability to recruit an mRNA to a specific nuclear location, we sought to test the impacts of nuclear speckle localization on splicing efficiency. To establish the baseline splicing efficiency and account for non-MCP dependent effects on GFP expression – including transfection or specific protein-dependent effects – we expressed each protein without MCP. We quantified the relationship between directed recruitment and splicing efficiency by measuring the difference in GFP fluorescence with and without MCP for each protein construct (ΔGFP) relative to protein levels (mCherry).

Recruitment of MS2-RNA specifically to speckle proteins SRRM1 or SRSF1 resulted in a non-linear change in splicing efficiency (ΔGFP) relative to protein levels (nonlinear four parameter logistic regression; R2 = 0.92 and 0.94, respectively; Figure 3F; Supplemental Figure 2A and 2B). To ensure that this observed effect is specifically due to nuclear speckle recruitment, we recruited this MS2-RNA to the diffusely localized splicing proteins SRSF3 and SRSF9 or to the nuclear lamina using LBR. In all cases, we observed that these conditions had no impact on splicing efficiency (Figure 3E-3F; Supplemental Figure 2C-2E).

These results indicate that direct recruitment of a pre-mRNA to nuclear speckle proteins, but not to other nuclear proteins, is sufficient to increase splicing efficiency. To ensure that this effect is specifically due to the ability of these proteins to localize within the nuclear speckle, we expressed a truncated form of SRRM1 that lacks the domain responsible for nuclear speckle localization but retains its catalytically active RNA processing domain20 (ΔNS-SRRM1; Figure 3G). We confirmed that ΔNS-SRRM1 no longer localizes within nuclear speckles (Figure 3G). Interestingly, expression of ΔNS-SRRM1 leads to a loss of the MCP-dependent increase in splicing efficiency and instead shows a response similar to that observed for other non-speckle-associated proteins (Figure 3H).

Together, these results demonstrate that directed recruitment of a pre-mRNA to nuclear speckles leads to a non-linear increase in mRNA splicing efficiency.

### Differential gene positioning around nuclear speckles corresponds to differential Pol II occupancy

Because gene organization around nuclear speckles impacts splicing efficiency, we sought to determine whether this organization changes between cell types. To explore this, we generated SPRITE maps in mouse myocytes derived from differentiated mm14 mouse myoblast cells. We computed genome-wide nuclear speckle distances from >14 million SPRITE clusters (Supplemental Figure 4A-4D) and observed that DNA regions located close to speckles correspond to genomic regions containing high-density of RNA Pol II in differentiated myocytes (Spearman correlation = 0.69, p<0.0001, Figure 4A-4B). Importantly, not all highly transcribed Pol II genes organize around the speckle; for example, the Chd2 gene on mouse chromosome 7 contains high levels of Pol II – comparable to that of the nearby Btbd1 gene – yet is located farther from the speckle, likely because Chd2 is transcribed from an otherwise Pol II sparse location (Figure 4A).

Next, we compared myocyte speckle distances to those measured in mouse ES cells. Overall, we observe that ~25% of the genome is speckle-proximal in either mouse ES or myocytes. Of these, ~40% are speckle proximal in both cell types whereas ~30% are speckle-proximal only in ESCs and the other ~30% specific to myocytes (Figure 4C). Because speckle proximity is correlated with Pol II density, we explored whether the changes in speckle proximity between myocytes and ES cells corresponded to changes in Pol II localization. Indeed, these unique speckle-proximal regions correspond to genomic regions that contain the largest differences in RNA Pol II between myocytes and ES cells (Spearman correlation = 0.52, p<0.001, Figure 4D). Similarly, genomic regions that are speckle-proximal in ES cells but not in myocytes correspond to regions that contain higher amounts of Pol II in ES cells relative to myocytes. For example, the genomic neighborhood containing the pluripotency marker Nanog is highly expressed and displays high speckle contact frequency in ES cells (Figure 4E-4F). In contrast, Nanog is not expressed in myocytes and the same genomic region displays a low speckle contact frequency (Figure 4F). Conversely, myogenic differentiation leads to widespread transcriptional upregulation of skeletal muscle specific genes, such as Titin (Ttn) and MyoD1 in myocytes. We observed that these regions were highly expressed and located proximal to nuclear speckle hubs in myocytes, whereas these same regions in mES cells were not expressed and were localized away from nuclear speckles (Figure 4E–5F; Supplemental Figure 4E).

**Figure 5:**
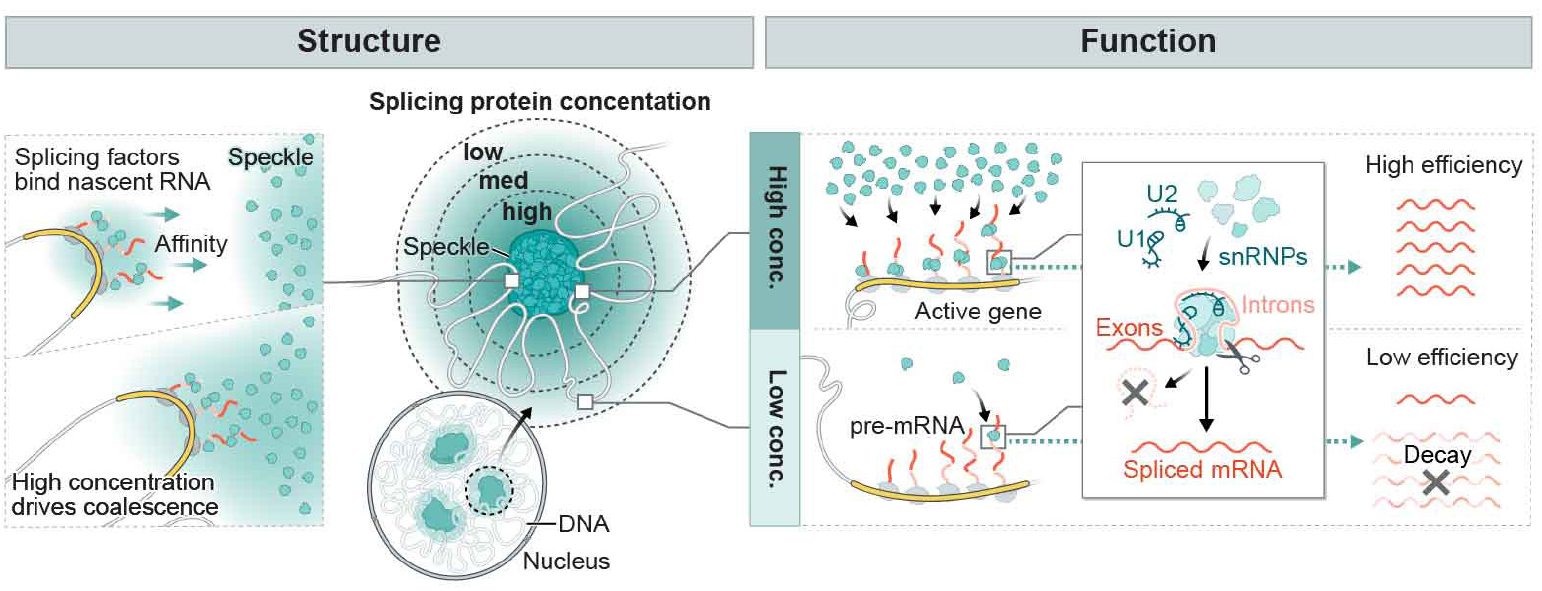
Integrated model for how gene organization around nuclear speckles impact splicing. Model of how 3D genome organization drives mRNA splicing. Because nascent pre-mRNAs have high affinity for splicing factors and Pol II dense regions contain the highest concentrations of nascent pre-mRNAs, these genomic regions can achieve multivalent contacts with splicing factors that are enriched within nuclear speckles. Because nuclear speckles contain the highest concentration of these factors within the nucleus, these multivalent contacts may drive coalescence (self-assembly) of these genomic DNA sites with the nuclear speckle. Genomic regions and pre-mRNAs close to nuclear speckles have higher levels of spliceosomes than regions farther away. Locally concentrating pre-mRNAs, genomic DNA, and spliceosomes at speckle-proximal regions leads to increased splicing efficiency whereas a speckle far gene transcribed at the same level is not spliced as efficiently.

To explore these changes more generally, we performed SPRITE on four distinct human cell types: H1 human embryonic stem cells (H1 hESC), H1 ES-derived endoderm cells, human foreskin fibroblasts (HFF-c6), and human lymphoblastoid cells (GM12878) (Figure 4G; Supplemental Figure 5A-C). Similar to the differential speckle contacts observed in mouse, we observed differential speckle localization for genes that are cell type specific. For example, the genomic region containing vimentin (also known as fibroblast intermediate filament) was most speckle proximal in HFF relative to the other three cell types. In contrast, the retinol binding protein 4 (RBP4) was most speckle proximal in ES-derived endoderm cells, consistent with findings that RPB4 is primarily expressed by endoderm-derived liver [76].

Together, these results reveal that distinct cell types display different speckle contacts, and these differences are associated with RNA Pol II density over genomic DNA regions (Figure 4H).

## DISCUSSION

Our results suggest a model that integrates the longstanding observations of nuclear speckles with the biochemistry of mRNA splicing. In this model, nuclear speckles consist of high concentrations of inactive spliceosomes which, when activated, diffuse away to engage pre-mRNAs [12,13,48,53,77]. When a nascent pre-mRNA is located closer to a speckle, there is a reduced volume through which the active spliceosomes need to diffuse to interact with the pre-mRNA. This decrease in diffusion volume creates a higher concentration of spliceosomes in the vicinity of speckle-close genes and thus results in increased spliceosome binding to these pre-mRNAs and conversion into spliced mRNA (Figure 5).

We demonstrate that genome organization around speckles differs by cell type, a finding consistent with recent observations [78,79]. Specifically, because speckle proximity is correlated with Pol II density and genes are differentially organized relative to speckles based on transcriptional activity, high levels of transcription may act to reposition genomic DNA closer to the nuclear speckle. Because nascent pre-mRNAs have high affinity for splicing factors and Pol II dense regions contain the highest concentrations of nascent pre-mRNAs, these genomic regions may achieve multivalent contacts with splicing factors that are enriched within nuclear speckles. These multivalent contacts may in turn drive coalescence (self-assembly) of these genomic DNA sites with the nuclear speckle [2] (Figure 5). Indeed, this self-assembly concept explains how newly transcribed ribosomal DNA genes and snRNA gene loci coalesce into the nucleolus [2,7] and Cajal bodies [17,80,81], respectively. Although RNA Pol II density is associated with speckle proximity [61], not all highly transcribed genes in a cell type are organized around the speckle. Because differential splicing efficiency would impact mRNA and protein levels in a cell, varying genome organization relative to speckles may drive differences in splicing efficiencies and therefore create another dimension of gene expression control.

mRNA splicing and Pol II transcription are known to be kinetically coupled [50,82–84] such that increasing the transcription of a gene leads to a non-linear increase in its splicing efficiency (referred to as ‘economy of scale’ splicing [85]). While individual splicing proteins have been shown to associate with the C-terminal domain of Pol II [83,86–91] direct binding of splicing factors to Pol II would predict a linear relationship between transcription and splicing and therefore cannot fully explain this coupling. Moreover, Pol II is not sufficient to stimulate splicing efficiency in cellular extracts [92]. This suggests that there must be some additional cellular mechanism required to functionally couple transcription and splicing in cells; our results suggest that this mechanism may be differential gene organization relative to nuclear speckles. Specifically, high levels of Pol II transcription would act to reposition genomic DNA into proximity with the nuclear speckle and increase splicing efficiency at these genes. Consistent with this notion, it was previously observed that economy of scale splicing also corresponds to an increased proximity between the gene locus and nuclear speckles [85]. Because the increase in spliceosome concentration achieved at DNA regions positioned at the nuclear speckle would exceed the proportional concentration of the pre-mRNAs transcribed at that locus, this model would explain the observed non-linear increase in splicing efficiency that is achieved when a gene is recruited to the nuclear speckle.

More generally, our results suggest a novel mechanism by which nuclear organization can coordinate regulatory processes in the nucleus and ensure robust non-linear control. Beyond speckles, there are many other bodies that similarly organize RNA processing enzymes with their co-transcriptional DNA and RNA targets [1,2,62]. These compartments include nascent ribosomal RNA loci and rRNA processing factors (e.g., snoRNAs, nucleolin) within the nucleolus [7,93], histone mRNAs and histone processing factors (e.g., U7 snRNA) in histone locus bodies [8,9], and snRNAs and their processing factors (e.g., scaRNAs) within Cajal bodies [10,11,94]. In each of these examples, these nuclear bodies organize around active transcription of the genes that they process [62]. Our results suggest that this structural arrangement may be an important and shared role for coordinating the co-transcriptional efficiency of RNA processing. Specifically, assembling genomic DNA encoding nascent pre-RNAs and their associated regulatory factors within the nucleus could act to increase the local concentration of these factors and therefore couple the efficiency of RNA processing to transcription of these specialized RNAs. This organization would enable localization of these RNA processing enzymes at their targets as they are being produced. The importance of ensuring robust and efficient co-transcriptional processing and coordinating these processes in space and time may explain why all known classes of RNA processing are associated with specialized nuclear bodies.

## ACKNOWLEDGEMENTS

We thank Allen Chen, Mackenzie Strehle, Drew Honson, Elizabeth Soehalim, Elizabeth Detmar, Say-Tar Goh, and Drew Perez for experimental help; Isabel Goronzy for computational assignment of speckle hubs; Lior Pachter, Tara Chari, Ben Riviere, Delaney Sullivan, and Noah Ollikainen for computational help; Michael Elowitz, Barbara Wold, Long Cai, Arjun Raj for reagents; Fangyuan Ding, Hong Yin, Joanna Jachowicz, Luke Frankiw, Yicheng Luo for helpful discussions; Igor Antoshechkin for sequencing; Aaron Lin for sequencing advice; Mackenzie Strehle, Drew Honson and Kent Leslie for critical comments on the manuscript; Job Dekker and Johan Gibcus for HFFc6 cell lines, Rene Maehr and Krishan Mohan Parsi for H1 ESC and H1 endoderm cell lines. Inna-Marie Strazhnik for illustrations and Shawna Hiley for editing. Imaging was performed in the Biological Imaging Facility with the support of the Caltech Beckman Institute and the Arnold and Mabel Beckman Foundation.

## AUTHOR CONTRIBUTIONS

P.B. and M.G. conceived the study, analyzed data, interpreted results, and wrote the manuscript. P.B., A.C., and M.G. designed experiments. P.B., A.C., O.E., S.A.Q., W.H., and M.R.B. performed experiments. B.E. performed RNA FISH and image analysis including nuclear segmentation and spot detection. Y.T analyzed seqFISH+ data. P.B. and M.G. supervised the work and M.G. acquired funding.

---

**Supplemental Figure 1:**
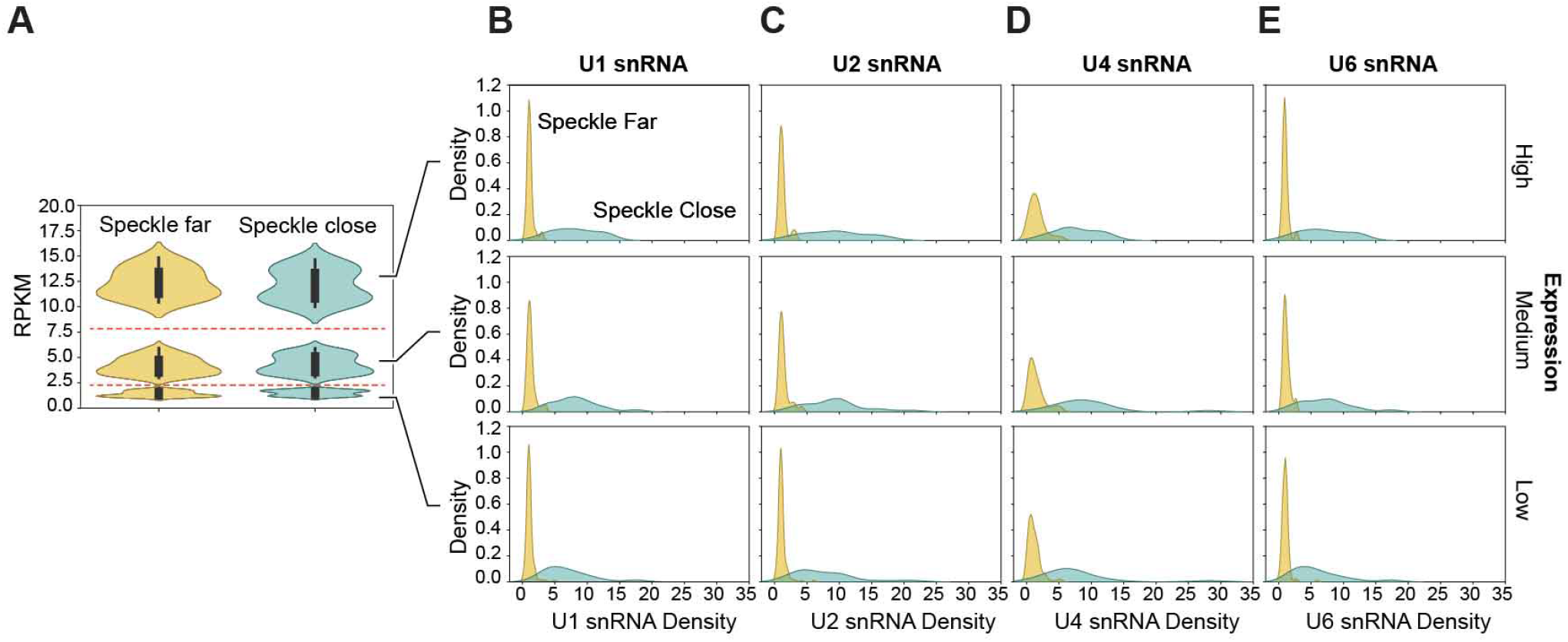
snRNA density for differently expressed genes. To ensure that splicing factor difference were not due to expression differences between speckle close and speckle far genes, we divided genes up based on expression ranges: high expression (RPKM > 10), medium expression (RPKM = 2.5-7.5), low expression (RPKM = 0-2.5). The distribution of expression within these ranges were the same for speckle close and speckle far genes. **(B)** U1 snRNA density is plotted for high (top), medium (middle), and low expression genes (bottom). **(C)** U2 snRNA density is plotted for high (top), medium (middle), and low expression genes (bottom). **(D)** U4 snRNA density is plotted for high (top), medium (middle), and low expression genes (bottom). **(E)** U6 snRNA density is plotted for high (top), medium (middle), and low expression genes (bottom).

**Supplemental Figure 2:**
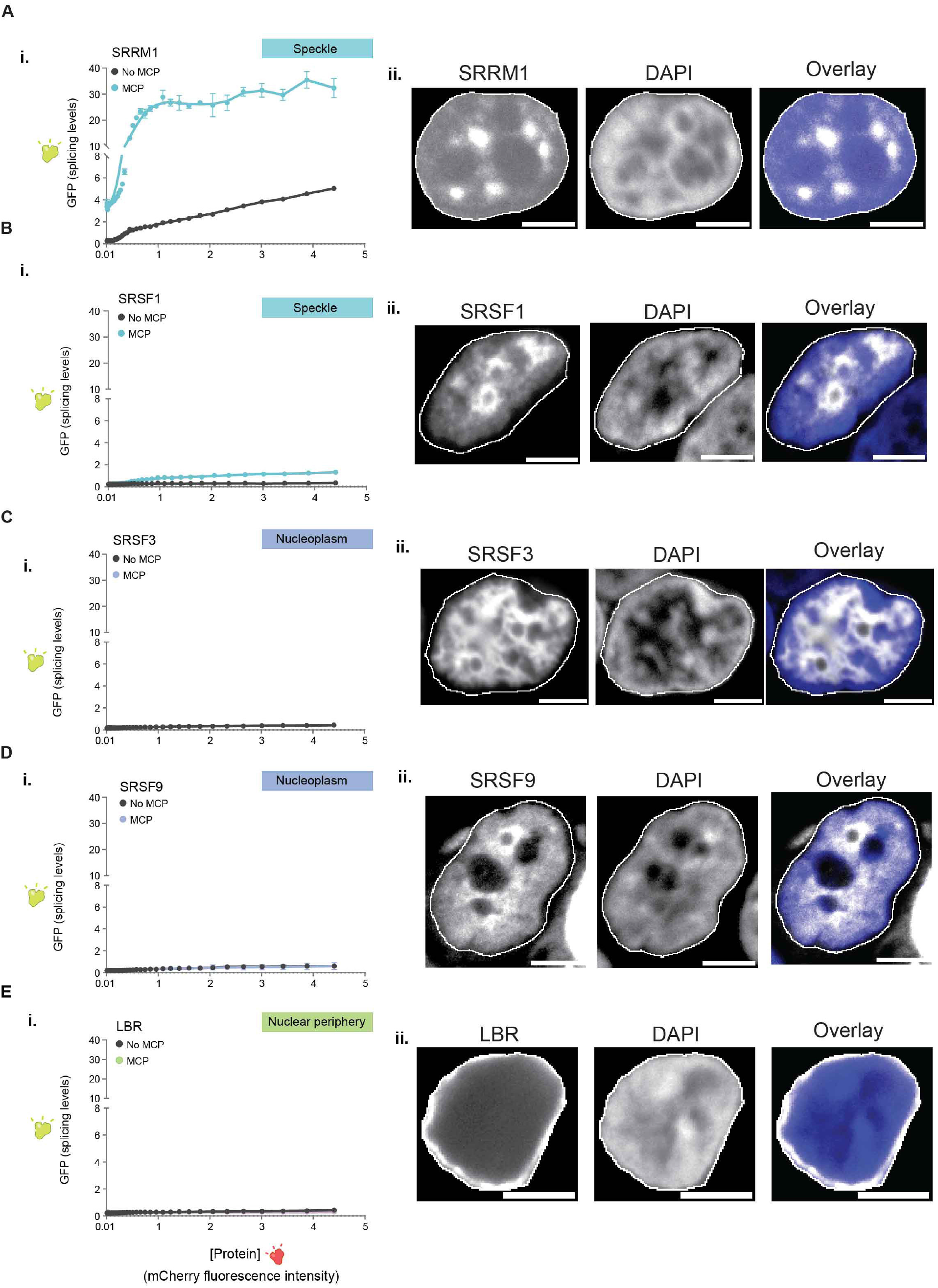
pre-mRNA organization around nuclear speckles drives splicing efficiency. (i) GFP fluorescence (splicing levels) (y axis) versus mCherry fluorescence intensity for constructs with MCP or without MCP (Left) for: **(A)** SRRM1, **(B)** SRSF1, **(C)** SRSF3, **(D)** SRSF9, **(E)** LBR. **(ii)** Imaging of each protein **(A-E)** with DAPI and overlay (Right) with nucleus outlined in white. Images are a more complete representation of those displayed in Figure 3B. Scale bars, 5 μm.

**Supplemental Figure 3:**
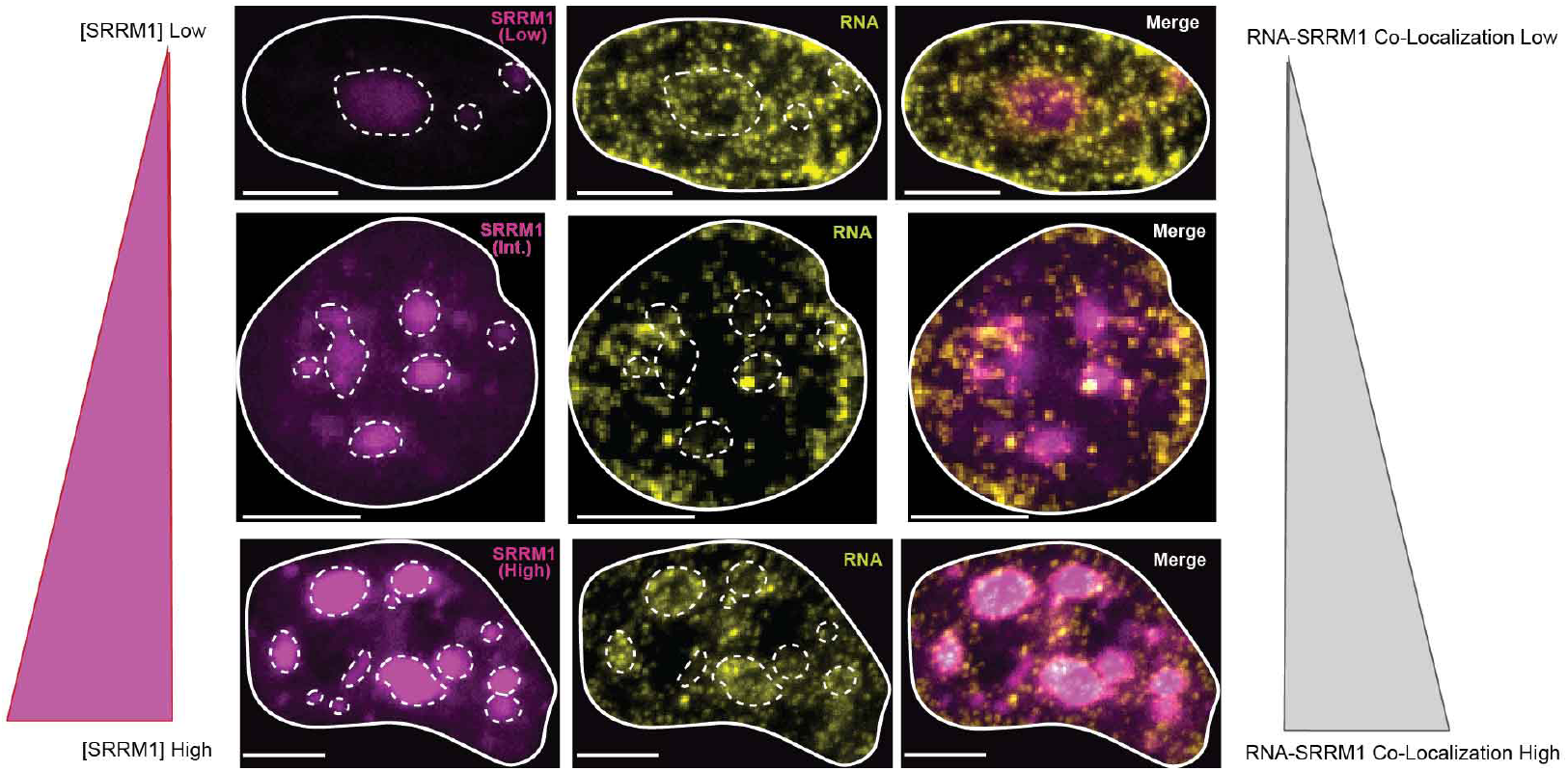
MS2-RNA localization to speckles is concentration dependent. SRRM1+MCP fluorescence microscopy (left) combined with RNA FISH (middle) and overlay (right). Low (top left), intermediate (middle left), and high (bottom left) SRRM1 expression. Scale bar is 5 μm.

**Supplemental Figure 4:**
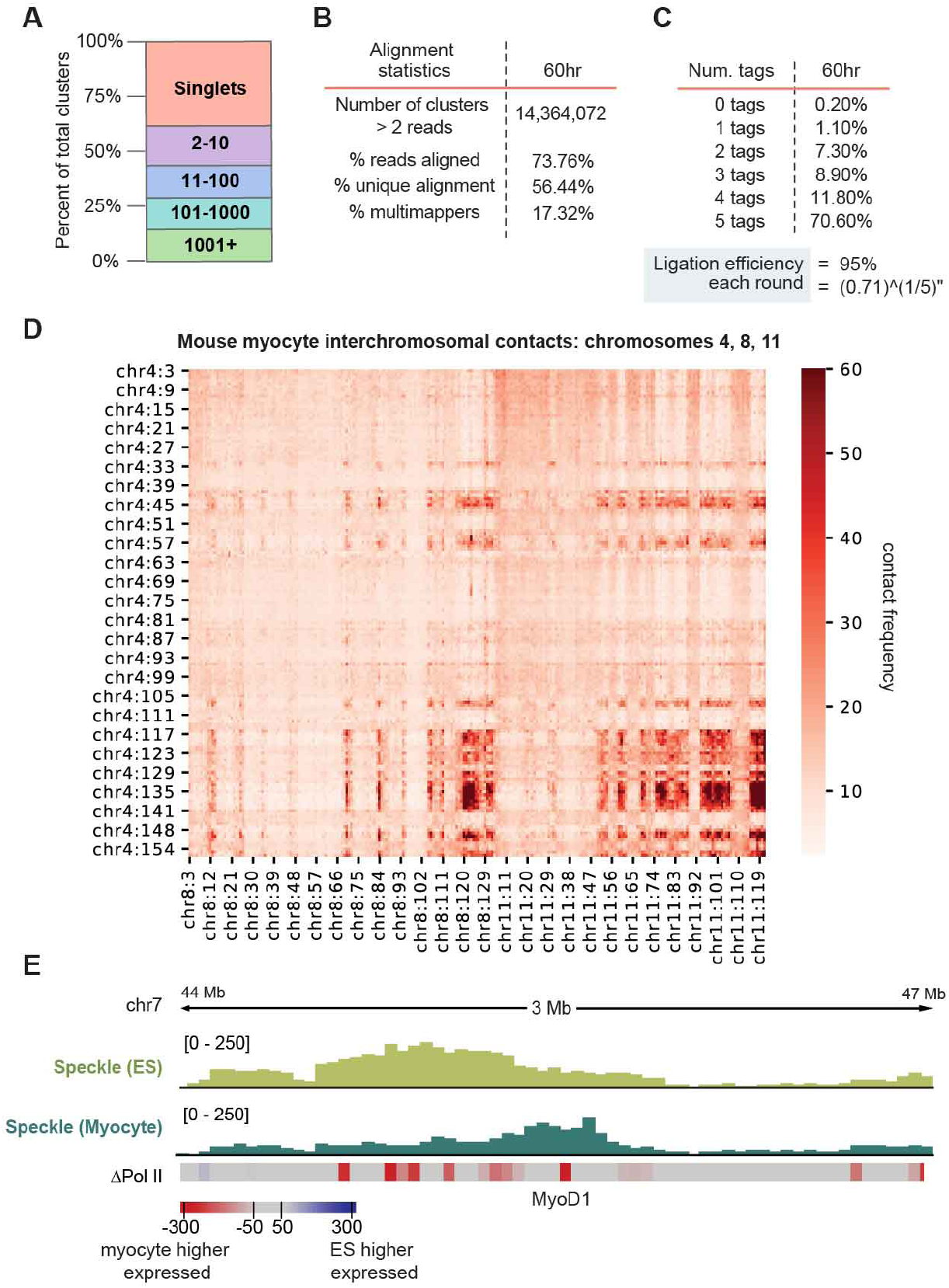
SPRITE analysis of mm14 myocyte cells. **(A)** Distribution of SPRITE cluster sizes for myocyte SPRITE. The percentage of reads was calculated for different SPRITE cluster sizes (1, 2-10, 11-100, 101-1000, and over 1001 reads) and reported as the percentage of total reads. Cluster size is defined as the number of reads with the same barcode. **(B)** Alignment statistics. **(C)** A summary of ligation efficiency statistics to confirm tags have successfully ligated to each DNA molecule. **(D)** Mouse myocyte interchromosomal contacts on chromosomes 4, 8, 11. **(E)** ES cell speckle contact frequency (light green) and skeletal muscle speckle contact frequency (dark green) for genomic locus near MyoD1 (expressed in myocyte). ΔPol II refers to difference in Ser2P-Pol II ChIP seq signal between mES cells and myocytes at 100-kb resolution, red is high in myocyte and blue high in ES.

**Supplemental Figure 5:**
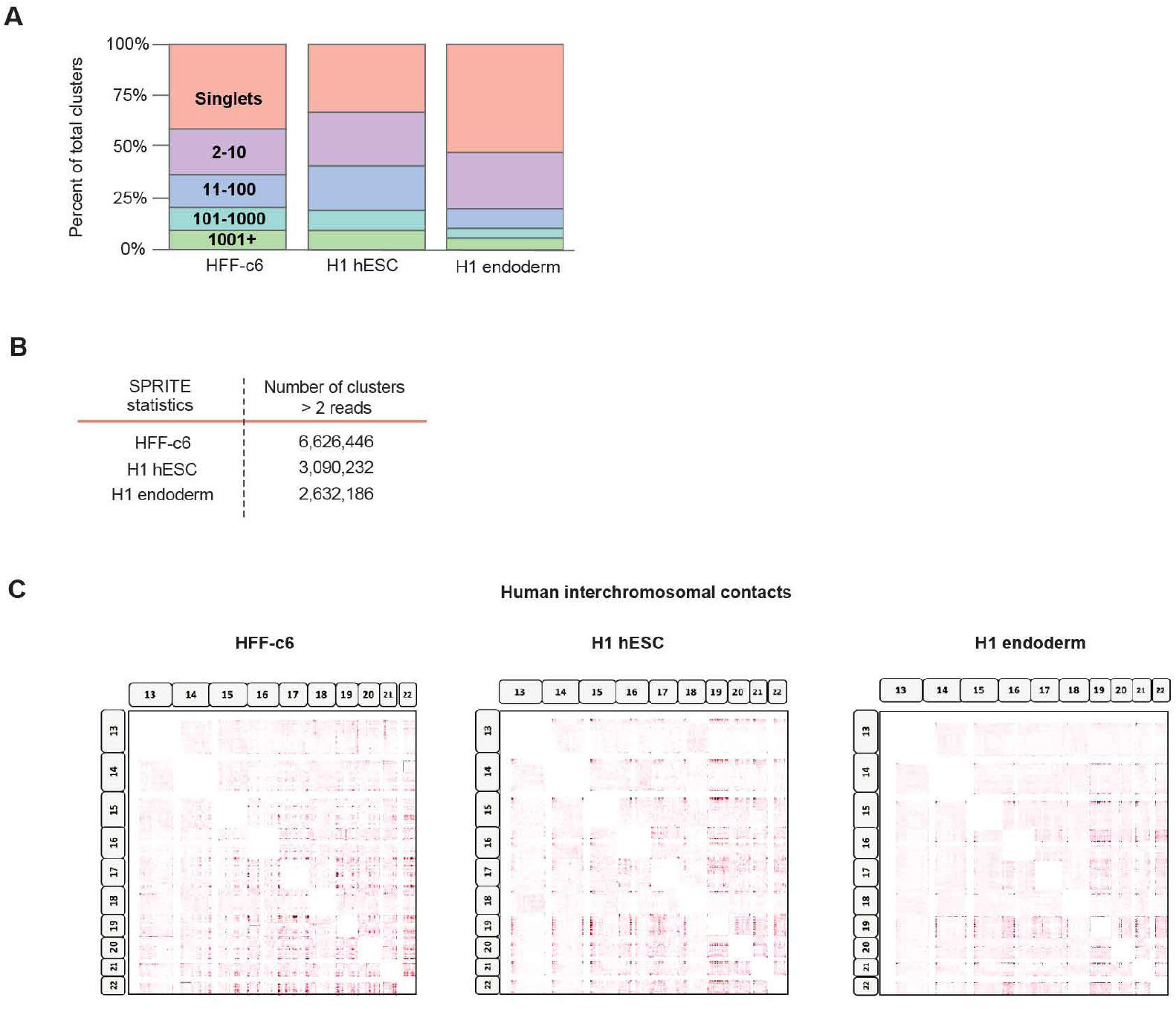
SPRITE analysis of human cells. **(A)** Distribution of SPRITE cluster sizes for HFF-c6, H1-hESC, and H1 endoderm SPRITE. The percentage of reads was calculated for different SPRITE cluster sizes (1, 2-10, 11-100, 101-1000, and over 1001 reads) and reported as the percentage of total reads. Cluster size is defined as the number of reads with the same barcode. **(B)** SPRITE statistics. **(C)** Human interchromosomal contacts on chromosomes 13 - 22.

## METHODS

### Visualization and Quantification of DNA seqFISH+ data

The DNA seqFISH+ and immunofluorescence data in mouse ES-E14 cells64 were downloaded from Zenodo (https://zenodo.org/record/3735329#.Y1t7Xuxuf0o). The pseudo-color images of DNA seqFISH+ spots were reconstructed from rounded voxel location of the decoded DNA seqFISH+ spots (seed values of 4 or 5), and then applied with a multidimensional Gaussian filter (sigma = 1) with scipy.ndimage.gaussian_filter package in python 3.7.13. The raw immunofluorescence images of nuclear speckles were reconstructed from csv files that contain intensity values of the SF3A66 antibody in each nucleus. The DNA seqFISH+ and immunofluorescence images were overlaid and contrasted by using ImageJ. The distance between the DNA locus and SF3A66 region was computed as previously described [64].

### Computing genome-wide speckle contact frequencies from SPRITE data

We computed genomic DNA distance to the speckle hub using the approach previously described [61]. Briefly, speckle hub regions were defined by clustering all significant inter-chromosomal contacts. We then computed a continuous distance metric for each bin of the genome (we state in figure legends and figures the resolution used: 1kb, 100kb, 1Mb) by identifying all SPRITE clusters containing both the genomic DNA bin and an inter-chromosomal DNA region contained within the speckle hub. We excluded all clusters where the genomic DNA bin and the speckle hub bin were only contained on the same chromosome in order to ensure that these distances were not driven by local (speckle-independent) contacts. Using these clusters, we then computed a contact score by divided by its respective cluster size (2/total number of reads within the cluster, as previously described [61]) and summed all contact scores for each genomic bin. This produces a continuous contact score for each genomic bin where low scores are farther from the speckle and higher scores are closer to the speckle.

### Comparison of SPRITE and SeqFISH+

To compare SPRITE and SeqFISH+Immunofluorescence measurements, we used SPRITE contact frequencies from contact maps binned at 1 Mb resolution focusing only on SPRITE clusters containing 2 to 1000 reads and down-weighting for cluster size (described above). The distance for SeqFISH+ represented the distance between the DNA spot and the periphery of the SF3A66 domain. When a DNA region and speckle are close, the SeqFISH+ distance is expected to be low and the SPRITE contact frequency is expected to be high.

### snRNA enrichment calculation from RNA & DNA SPRITE

We computed RNA-DNA contacts frequencies for U1, U2, U4, and U6 snRNAs in 1Mb bins across the genome, weighted by cluster size. For the same 1-Mb bins, we computed each genomic bins’ speckle hub contact frequency. To calculate transcription rate, we labeled mES cells for 10 minutes with 5-ethynyl uridine and purified the resulting RNA as previously described [95]. We aligned reads to mm10 and calculated reads per kilobase of gene per million reads (RPKM) mapped for each gene. After this, we computed the median RPKM for the region and filtered bins by expression level (RPKM 1-2 (low), 3-6 (medium), and 10-15 (high).

To compute snRNA enrichment genome-wide, we first computed speckle contact frequencies of each genomic bin at 1 Mb. Next, we computed the contact frequency of U1, U2, U4, and U6 snRNAs for each of these same 1 Mb bins. We then performed a rank normalization of speckle contacts and defined speckle far as the regions corresponding to the lowest 5% of speckle contact frequencies and speckle close as the top 5% of regions. To normalize all values to the same range, the contact score of each snRNA bin value was divided by the median of the speckle far contacts. To compare only regions of equivalent expression, we thresholded regions corresponding to low, medium or high expression. To do this, we computed the median RPKM for the region and filtered bins by expression level (RPKM 1-2 (low), 3-6 (medium), and 10-15 (high). Density plots for speckle close and speckle far regions, for each snRNA, and for each expression level were plotted using the seaborn kde function.

### U1 snRNA enrichment calculation from psoralen crosslinking (RAP-RNA AMT)

To compute direct U1 snRNA-pre-mRNA interactions, data from RAP-RNA from AMT crosslinking [67] (GEO IDs: GSM1348350 (input RNA AMT) and GSM1348348 (U1 AMT RAP RNA)) was re-analyzed. In this procedure, cells are treated with a psoralen crosslinker to form direct crosslinks between directly base pair hybridized RNA-RNA sequences. Affinity capture for U1 snRNA and sequencing of associated RNAs identifies the RNAs that were directly bound to U1. To normalize for transcript abundance, input RNA libraries were sequenced in parallel.

The enrichment of U1 snRNA on pre-mRNAs was computed by dividing the contact frequency for each 100-kb genomic bin in the capture by the input. Speckle contact frequencies for the same 100-kb bins were computed as above. Also as above, we performed a rank normalization of speckle contacts and defined the top and bottom 5% of speckle contact frequencies as speckle close and speckle far. We plotted the density for all speckle close and far regions for the U1 snRNA using the seaborn kde function.

### Nascent splicing efficiency calculation from chromatin RNA sequencing

Total chromatin RNA sequencing [96] was re-analyzed from GEO ID: GSM2123095 and re-aligned using the kallisto-bustools workflow [97] to two references separately: a cDNA reference (for exon reads and exon-exon junction reads) and a genomic DNA reference genome (for exon-intron and intron reads). Splicing ratio was computed as the fraction of normalized exon counts over normalized intron + exon (total) counts. We filtered for speckle close and far regions as above and plotted the distribution of percent splicing using the seaborn kde function. For the continuous distribution plot, we plotted all speckle contact frequencies (x-axis) versus the average splicing ratio in each of 50 bins, where each bin contains at least 20 genes.

### Splicing efficiency calculation from RNA & DNA SPRITE (RD-SPRITE)

Because RNA & DNA SPRITE captures interactions occurring between DNA and RNA, we reasoned that any mRNA that was in a SPRITE cluster with its own DNA locus corresponded to nascent chromatin associated RNA. Indeed, we previously showed that this approach accurately captures and quantifies nascent pre-mRNA levels [70]. Using these clusters, we computed splicing efficiency based on the total number of exon reads in a nascent genomic bin divided by the total number of exons and introns (total pre-mRNA reads) within that same bin. To ensure that we had robust coverage to estimate this frequency, we filtered for genomic regions that contained at least 50 RNA reads (exons + introns). We filtered for speckle close and far regions as above and plotted the distribution of percent splicing using the seaborn kde function. For the continuous distribution plot, we plotted all speckle contact frequencies (x-axis) versus the average splicing ratio in each of 50 bins, where each bin contains at least 20 genes.

### Plasmid generation for MS2-MCP assay

#### mCherry-fused, MCP-tagged expression plasmid

The Gateway destination plasmid pCAG-NSTF-DEST-V5 (gift from P. McDonel) was modified by digestion/ligation methods to add mCherry in frame following the V5 tag. This was the -MCP destination vector. To generate the +MCP version, digestion/ligations methods were used to remove the NSTF cassette and to replace it with 2xMCP. These destination vectors were used in Gateway LR recombination reactions with entry clones for each protein of interest. Entry clones were obtained from DNASU.

#### deltaIDR-SRRM1 entry clone

The SRRM1 entry clone from DNASU was modified using the Q5 site directed mutagenesis kit (New England Biolabs) to delete the predicted disordered region as annotated by Uniprot (https://www.uniprot.org/uniprotkb/Q8IYB3/entry). The resulting clone lacked one additional amino acid at the C-terminus as determined by sanger sequencing of the clone and alignment with the predicted sequence.

### Imaging analysis for MS2/MCP reporter assay

#### RNA FISH

To visualize RNA localization in MCP-MS2 recruitment assays, we co-transfected HEK293 cells with splicing reporter and domain recruitment constructs then performed single-molecule RNA FISH as previously described [98]. 24-hours after transfection, we rinsed samples once with 1X PBS then fixed in 4% buffered formaldehyde for 10 minutes at room temperature. Following fixation, we rinsed the samples twice with 1X PBS then permeabilized in 70% ethanol overnight at 4°C. For hybridization, we rinsed the samples once with wash buffer (10% formamide 2X SSC) then added hybridization buffer (10% formamide, 10% dextran sulfate, 2X SSC) containing RNA FISH probes targeting GFP RNA. These probes were kindly provided by Arjun Raj (University of Pennsylvania). After adding the hybridization solution, we covered samples with glass coverslips and hybridized overnight at 37°C in a humidified container. Following hybridization, we rinsed the samples once with wash buffer to remove coverslips and then washed twice for 30 minutes at 37°C. We added 50 μg/mL 4’,6-diamidino-2-phenylindole (DAPI) to the second wash to stain nuclei. Following washes, we rinsed the samples twice with 2X SSC, added SlowFade™ Diamond Antifade solution and proceeded with imaging on a Nikon spinning-disk confocal equipped with Andor Zyla 4.2P sCMOS camera, Nikon LUNF-XL laser unit, and Yokogawa CSU-W1 with 50 μm disk patterns. For each sample, we selected at least ten positions on the basis of DAPI signal and acquired z-stacks at 0.5 μm intervals using a x60 oil objective.

### Immunofluorescence

Cells were fixed on coverslips with 4% formaldehyde in PBS for 15 min at room temperature and permeabilized with 0.5% Triton X-100 in PBS for 10 min at room temperature. After washing twice with PBS containing 0.05% Tween (PBSt) and blocking with 2% BSA in PBSt for 30 min, cells were incubated with primary antibodies for anti-SC35 antibody at 1:200 dilution (Abcam, ab11826) overnight at 4 °C in 1% BSA in PBSt. After overnight incubation at 4 °C, cells were washed three times in 1× PBSt and incubated for 1 h at room temperature with secondary antibodies labeled with Alexa fluorophores (Invitrogen) diluted in 1× PBSt (1:500). Next, coverslips were washed three times in PBSt, rinsed in PBS, rinsed in double-distilled H2O, mounted with ProLong Gold with DAPI (Invitrogen, P36935) and stored at 4 °C until acquisition.

### Image analysis

To quantify RNA recruitment to nuclear lamina or speckles, we used Cellpose (https://github.com/mouseland/cellpose) to segment nuclear boundaries based on DAPI signal and used the Raj Lab smFISH pipeline (https://github.com/arjunrajlaboratory/rajlabimagetools) to localize intranuclear reporter RNA [98,99]. We then quantified mCherry fluorescence intensity at the position of each reporter RNA molecule. To account for heterogeneity in mCherry expression across cells, we calculated the rank pixel intensity to measure relative RNA-mCherry colocalization across conditions. We note that expression heterogeneity precluded us from segmenting speckle domains consistently across cells. In addition, to account for heterogeneity in co-transfection efficiency, we had a blinded author manually select non-mitotic cells co-transfected with both the splicing reporter and the domain recruitment construct.

Due to the sequence and length (GUACAUCUGGUCCAUCCUUCCUAGCUGCGUCCUGGUGGCGC AGGUGUGG GGGAUCGGCAGGUGCCUACCACUAUGCUGUCUAUUACAG; 88 nucleotides;) the intron in our splicing reporter our splicing reporter, we were unable to design smFISH probes selectively targeting nascent RNA. Instead, we used a probeset targeting exons present in both nascent and mature RNA. Since only nascent (unspliced) RNA contain the MS2 hairpin, our results likely underestimate the extent of reporter RNA recruitment.

### Overexpression of MS2/MCP constructs in HEK293T

For MS2/MCP experiments that required a wide range of protein expression, human HEK293T cells were used instead of mESCs because they allow for a wide range of expression levels and enabled investigation of the effect of varying concentrations of proteins (with and without recruitment) on splicing efficiency.

HEK293T cells were cultured in complete media consisting of DMEM (GIBCO, Thermo Fisher Scientific) supplemented with 10% FBS (Seradigm Premium Grade HI FBS, VWR), 1X penicillin-streptomycin (GIBCO, Thermo Fisher Scientific), 1X MEM non-essential amino acids (GIBCO, Thermo Fisher Scientific), 1 mM sodium pyruvate (GIBCO, Thermo Fisher Scientific) and maintained at 37°C under 5% CO2. For maintenance, 800,000 cells were seeded into 10 mL of complete media every 3-4 days in 10 cm dishes.

To assess splicing efficiency of the MS2 splicing reporter, exons 5-6 of mouse IRF7 (ENMUST00000026571.10) containing its endogenous intron were fused upstream of 2A self-cleaving peptide and eGFP and cloned into an MSCV vector (PIG, Addgene) [100]. This splicing reporter has a stop codon embedded within the intron, thereby only when the reporter is spliced will eGFP be translated. An MS2 stem loop was introduced into the intron to enable recruitment of the nascent pre-mRNA splicing reporter specifically to MCP-tagged proteins. The MS2 and tagged protein constructs were co-transfected into HEK293Ts. Splicing, as measured by GFP fluorescence, was assayed 24 and 48 hours after transfection by flow cytometry (Macsquant) and analyzed using FloJo analysis software. Transfections were performed using BioT transfection reagent (Bioland) according to the manufacturer’s recommendations. Transfected constructs included SRRM1, SRSF1, SRSF3, SRSF9, and LBR; all constructs were fused to a C terminal mCherry tag. Constructs harboring the MCP tag were fused to two tandem repeats of the MCP peptide at the N terminus.

### GFP expression as a function of various proteins fused to mCherry

For each construct (+/− MCP), we sorted on GFP and mCherry (doubly transfected cells). Because each protein expressed is fused to an mCherry, we assumed that the increase in mCherry fluorescence is proportional to the concentration of the protein of interest within the cell. As a control, we also sorted cells that contained constructs expressing GFP only or mCherry only to ensure there was no spillover of the fluorescence detection between constructs. Additionally, we sorted untransfected cells to set a baseline threshold to filter out cells with background autofluorescence. To that end, because the range of expression is variable due to differences in transfection efficiency/etc, we thresholded cells that contained the same range of mCherry fluorescence intensity (between 0 and 5) and contained non-zero GFP values. The upper threshold of 5 for mCherry fluorescence was chosen because that represents the upper bound of mCherry expression for the protein construct with the overall lowest levels of expression (SRRM1 + MCP). Next, because most points for all constructs were in the lower range of mCherry fluorescence (0-1), mCherry fluorescence (x axis) for each construct was logarithmically binned to 50 bins between 0-5 and the average GFP value for each bin was plotted. Each construct had at least three replicates. Data were merged after binning and the S.E.M. is plotted.

### Difference in GFP expression calculations

For each x-value (50 mCherry bins), the difference in average GFP fluorescence was computed between MCP and no MCP constructs. The average difference of at least three replicates were plotted for all constructs with S.E.M.

### Non-linear regression statistics

Data from each construct (ΔGFP for SRRM1, SRSF1, SRSF3, SRSF9, and LBR) were fitted using a four-parameter logistic curve and goodness of fit was calculated using GraphPad Prism 9 software.

### Myoblast cell culture and differentiation

C2C12 mouse skeletal myoblasts were passaged at 50-60% confluency every 1-2 days using the Wold lab protocol: (https://www.encodeproject.org/documents/a5f5c35a-cdda-4a45-9742-22e69ff50c9c/@@download/attachment/C2C12_Wold_protocol.pdf). Undifferentiated myoblasts grow in growth medium (20% fetal bovine serum). Myogenic differentiation was initiated upon reaching confluence by switching the cells to medium containing 2% horse serum supplemented with insulin. Differentiation was performed for 60 hours by rinsing fully confluent cells once with PBS and adding 25mL of low-serum differentiation medium. Fresh differentiation medium was changed every 24 hours up to the 48h timepoint and 12 hours afterward were crosslinked using SPRITE crosslinking procedures60.

### Human cell culture

HFFc6 cells were cultured in DMEM medium supplemented with 20% heat-inactivated FBS. Cells were crosslinked according to our previous SPRITE crosslinking procedure60. Details of culture conditions are available on the 4DN portal

https://data.4dnucleome.org/biosources/4DNSRC6ZVYVP/.

H1 hESC cells were maintained on matrigel matrix (Corning, 354277) in feeder free media using mTeSR1 (Stemcell Tech, 85850). Every 4-5 days cells were passaged using ReLeSR reagent (Stemcell tech, 05872).

H1 hESC differentiation to Definitive Endoderm

A detailed protocol from Maéhr lab is available on the 4DN portal

(https://data.4dnucleome.org/protocols/680ed3dd-04aa-49bc-aac0-8c88da6fddb6/).

Briefly, H1 hESC cells were grown to 80-90% confluency, dissociated into single cells, pelleted and resuspended in mTeSR1 supplemented with 1uM Y27632 (Tocris, 1254). Cells were seeded onto a 6 well coated plate with Growth Factor Reduced Matrigel. On day 1, cells were fed with mTeSR1 and incubated for 24hrs. On day 2, cells were changed with fresh media containing RMPI1640 (Thermo, 21870) supplemented with 0.2% Hyclone FBS (GE Healthcare, SH30070.03) 100 ng/mL Activin A (R&D Systems, 338-AC-01M), 3 μM CHIR 99021 (Tocris, 4423), and 50 nM PI 103 (Tocris, 2930) and incubated for 24hrs. On day 3, cells were changed with fresh media containing RPMI1640 supplemented with 0.2% Hyclone FBS, 100 ng/mL Activin A, and 250 nM LDN-193189 (Tocris, 6053) and incubated for 24hrs. On day 4, Cells were changed again with fresh media containing RPMI1640 supplemented with 0.2% Hyclone FBS, 100 ng/mL Activin A, and 250 nM LDN-193189 (Tocris, 6053). On day 5, cells were crosslinked with SPRITE crosslinking procedures as previously described.

### SPRITE cluster size calculations

DNA SPRITE and RNA & DNA-SPRITE were performed as previously described62. Unless stated otherwise, all analyses were based on SPRITE clusters of size 2–1000 reads. These cluster sizes were chosen to be consistent with the analysis in our previous papers, where we showed that many known structures such as TADs, compartments, RNA-DNA and RNA-RNA interactions, etc., occur within SPRITE clusters containing 2–1000 reads. GM12878 SPRITE data was generated previously61.

### Speckle hub definition

We computed speckle hub contacts from the myocyte data and human cells using the same approaches as previously described61. Briefly, the speckle hubs were defined by computing all inter-chromosomal contacts from DNA SPRITE at 1Mb resolution. Using these contacts, we computed p-values to identify all significant inter-chromosomal contacts and clustered these regions. As observed in the mouse ES data, we identified two, mutually exclusive sets of DNA regions, one of these two sets corresponds to the speckle hub and the other being the nucleolar hub. We used this speckle hub region to compute the speckle distance for each region of the genome by computing the number of SPRITE clusters containing the genomic DNA region and at least one of the regions contained within the nuclear speckle hub. To exclude this calculation being dominated by linearly proximal contacts on the same chromosome, we only counted clusters if they contained the genomic region of interest and a speckle hub region that was not contained on the same chromosome. Overall, the distribution of speckle hub scores across 1 megabase genomic regions are similar between mouse ES cells and myocytes, although the precise regions differ (see “SPRITE speckle hubs contact frequency” section).

### Comparing SPRITE datasets

To map and compare speckle hub contact frequencies (mouse ES vs myocyte; human SPRITE datasets) in each cell type, we performed a quantile normalization of the speckle hub contacts for each cell line to account for differences in coverage for each SPRITE.

